# Epithelial morphogenesis in the perinatal mouse uterus

**DOI:** 10.1101/2020.01.28.922385

**Authors:** Zer Vue, Richard R. Behringer

## Abstract

**Background:** The uterus is the location where multiple events occur that are required for the start of new life in mammals. The adult uterus contains endometrial or uterine glands that are essential for female fertility. In the mouse, uterine glands are located in the lateral and anti-mesometrial regions of the uterine horn. Previous 3D-imaging of the adult uterus, its glands, and implanting embryos has been performed by multiple groups, using fluorescent microscopy. Adenogenesis, the formation of uterine glands, initiates after birth. Recently, we created a 3D-staging system of mouse uterine gland development at postnatal time points, using light sheet fluorescent microscopy. Here, using a similar approach, we examine the morphological changes in the epithelium of the perinatal mouse uterus.

**Results:** The uterine epithelium exhibits mesometrial-antimesometrial (dorsoventral) patterning as early as three days after birth (P3), marked by the presence of the mesometrially-positioned developing uterine rail. Uterine gland buds are present beginning at P4. Novel morphological epithelial structures, including a ventral ridge and uterine segments were identified.

**Conclusions:** The perinatal mouse uterine luminal epithelium develops mesometrial-antimesometrial (dorsal-ventral) morphologies at 3-4 days post-partum. Between 5-6 days post-partum uterine epithelial folds form, defining alternating left-right segments.

**Bullet points:**

- Morphological patterning events in the perinatal uterine epithelium are not well described.
- Light sheet microscopy was used to generate volumetric reconstructions of the perinatal mouse uterine epithelium.
- At postnatal day 3 (P3), the uterine epithelium shows the first signs of dorsoventral pattern, with the presence of the forming mesometrially-positioned uterine rail.
- The first morphological indication of uterine adenogenesis begins at P4.
- Novel morphological structures were identified from volumetric reconstructions, including the presence of a ventral ridge (another sign of dorsoventral pattern) and uterine segmentation.

## Introduction

The mammalian female reproductive tract organs, including the oviducts, uterus, and vagina, are essential for the generation of progeny and a frequent site of human disease, including infertility, fetal loss, and cancer. The uterine lumen is lined by an epithelium called the luminal epithelium. Endometrial or uterine glands extend from the luminal epithelium into the surrounding stroma that together make up the endometrium. The stroma is surrounded by the two smooth muscle layers of the myometrium.

Uterine glands are important for survival and development of the mammalian embryo (Gray, Bartol et al. 2001). The primary technique used to visualize uterine glands is tissue sectioning and histology. In two-dimensional (2D) histological sections, uterine gland tissue is located within the stroma and is observed as discrete round or oval epithelial units, typically not continuous with the luminal epithelium (Filant, Zhou et al. 2012). In the past, this 2D histological method was used to generate models to describe how uterine glands develop. At postnatal day (P) 0 (day of birth), the mouse uterine horn consists of a central lumen lined by a simple epithelium, surrounded by undifferentiated mesenchyme cells. No uterine glands are present. In the mouse, between birth and P5, epithelial invaginations (buds) appear, indicative of the forming glandular epithelium (Kurita, Cooke et al. 2001, Cooke, Ekman et al. 2012). In addition, during this period, the mesenchymal layers of the uterus become radially oriented and distinct: the endometrial stroma and the inner circular myometrium (Brody and Cunha 1989). These initial epithelial buds do not become histologically distinct as uterine glandular tissue until P7 (Branham, Sheehan et al. 1985). Nascent glands elongate into the stroma by P10, while the outer longitudinal layer of the myometrium becomes organized into bundles (Brody and Cunha 1989). By P15, the basic adult configuration of the mouse uterus is established (Hu, Gray et al. 2004). Histological cross sections of the adult mouse uterine horn show mesometrial-antimesometrial (dorsal-ventral) differences. For example, uterine glands are located in the lateral and anti-mesometrial regions of the stroma (Hondo et al. 2007).

Recently, new imaging technologies have significantly facilitated the visualization of cells and tissues. This has resulted in the generation of high-resolution three-dimensional images of fluorescently-labeled reproductive tract tissues (Arora, Fries et al. 2016, Belle, Godefroy et al. 2017, Vue, Gonzalez et al. 2018, Yuan, Deng et al. 2018). These new imaging techniques have been used to determine uterine gland morphologies in three dimensions (Arora, Fries et al. 2016, Vue, Gonzalez et al. 2018, Yuan, Deng et al. 2018). Taking advantage of methods we developed, we generated 3D reconstructions of the postnatal uterine epithelium, revealing the structure of developing uterine glands. These 3D reconstructions supported the conclusion that no uterine glands are located at the mesometrial side of the uterine horn where the broad ligament is present. Interestingly, the 3D morphology of the uterine epithelium in this aglandular region revealed a structure we termed the “uterine rail” (Vue et al. 2018). The lack of glands in the mesometrially-located stroma and the presence of the uterine rail indicate epithelial patterning.

Here, we investigate the initial morphological changes and asymmetries that form in the epithelium of the perinatal mouse uterus, using light sheet fluorescent microscopy and 3D reconstructions. We find that the uterine rail forms by P3, indicating the development of mesometrial-antimesometrial (dorsal-ventral) polarity. By P4, Stage 1 uterine gland buds are found invading into the stromal compartment, marking the initiation of adenogenesis. At P4, we discovered the presence of a ventral ridge that forms anti-mesometrially in the epithelium. Subsequently at P6, two rows of uterine gland buds form on the ventral ridge. At P6, uterine folds begin to form that extend laterally into the stroma in a repetitive pattern creating segments. These morphologies identify epithelial patterning events in the uterine epithelium, resulting in dorsoventral patterning and anteroposterior segmentation, within the first week after birth.

## Results

### 3D reconstructions of the epithelium within the perinatal mouse uterus

At birth, a simple epithelium lines the lumen of the uterus, forming a tube lacking uterine glands (Vue et al., 2018). The morphological changes that occur in the uterine epithelium soon after birth are not well described. Indeed, there is very little volumetric information to deduce structures in three dimensions. To investigate the morphological changes that occur in the uterine epithelium during the perinatal period, we performed whole-mount immunofluorescent staining of uterine horns at P3, P4, P5, P6, P7 and P8, using the TROMA-1 (cytokeratin 8/18) antibody (**Fig. 1**). Previously, the TROMA-1 antibody has been shown to label the epithelial cells in the uterus (Brulet, Babinet et al. 1980). Using light sheet fluorescent microscopy, we were able to acquire volumetric images of the epithelium in the uterus, revealing epithelial structures (**Fig. 1A, E, I, M, Q, U**). Using Imaris software, we reconstructed 3D opaque surface renderings of the uterine epithelium (**Fig. 1B, F, J, N, R, V**). In addition, we were able to view the epithelium at higher magnifications (**Fig. 1C, G, K, O, S, W**) and in optical cross sections (**Fig. 1D, H, L, P, T, X**). These 3D reconstructions for P3-P8 uterine epithelium are presented in **Movies S1-S6**. From this 3D imaging of the uterine epithelium, we investigated the timing of uterine gland formation and have discovered novel morphological structures that have not been appreciated previously using traditional two dimensional histological approaches.

**Fig. 1.**
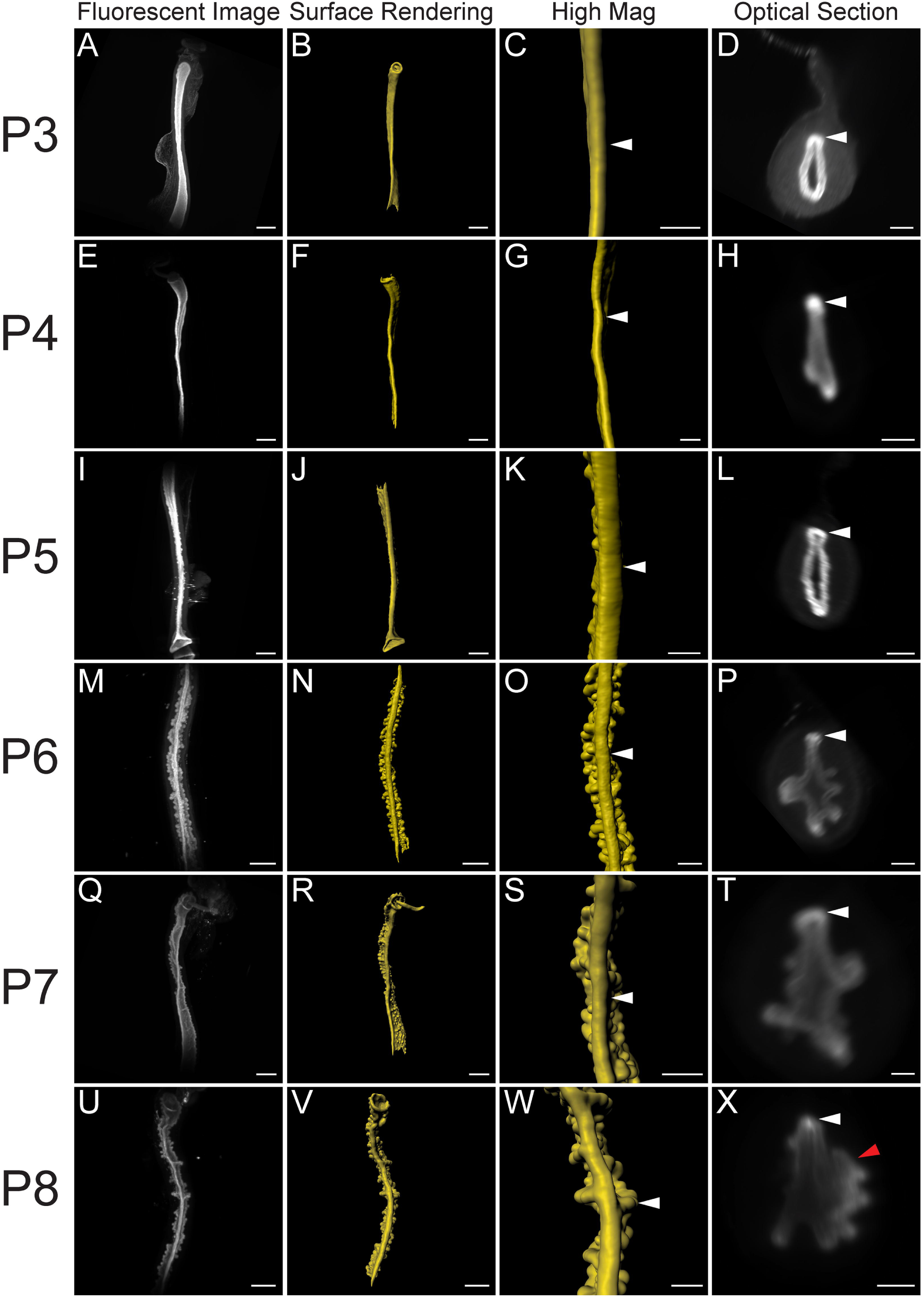
Volumetric image analysis of the perinatal mouse uterus. Fluorescent light sheet images and 3D reconstructions of the perinatal mouse uterus. Mesometrial view. (**A, E, I, M, Q, U**) TROMA-1 stained immunofluorescent images of the uterine epithelium. Mesometrial view. (**B, F, J, N, R, V**) Pseudo-colored surface renderings of the uterine epithelium corresponding to the first column. Mesometrial view. (**C, G, K, O, S, W**). High magnification surface renderings of the uterine epithelium corresponding to the second column. (**D, H, L, P, T, X**) Optical cross-sections of fluorescent light sheet images of TROMA-1 stained uteri. Mesometrium to the top. White arrowheads correspond to the uterine rail in the third and fourth columns. Red arrowhead corresponds to a lateral epithelial fold. (**A-D**) P3, (**E-H**) P4, (**I-L**) P5, (**M-P**) P6, (**Q-T**) P7, and (**U-X**) P8. Scale bars for (**A-B, E-F, I-J, M-N, Q-R, U-V**) = 500 μm. Scale bars for (**C-D, G-H, K-L, O-P, S-T, W-X**) = 200 μm.

### Initiation of uterine rail formation, an indication of mesometrial-antimesometrial (dorsoventral) patterning

We previously discovered the uterine rail by 3D imaging of the uterine epithelium at P8, P11, and P21 (Vue, Gonzalez et al. 2018). This epithelial structure is aglandular and runs along the mesometrial pole of each uterine horn. The presence of the uterine rail provides morphological evidence for mesometrial-antimesometrial (dorsoventral) patterning in the uterine epithelium. To determine when the uterine rail forms, we examined stages before P8. Consistent with previous data (Vue, Gonzalez et al. 2018), we found that the uterine rail is not present at P0 (**Fig. 2A, B**). However, the rail is present at P3 (**Fig. 2C, D**, n=6). The uterine rail appears fully formed and distinct by P8 (**Fig. 2E, F**, n=8). Single cross-section images of the z-stack from P3, P5, and P8 show that the uterine rail has a horseshoe appearance (**Fig. 2D, F**) that is not present at P0 (**Fig. 2B**). These findings suggest that the uterine epithelium develops dorsoventral polarity between P0 and P3.

**Fig. 2.**
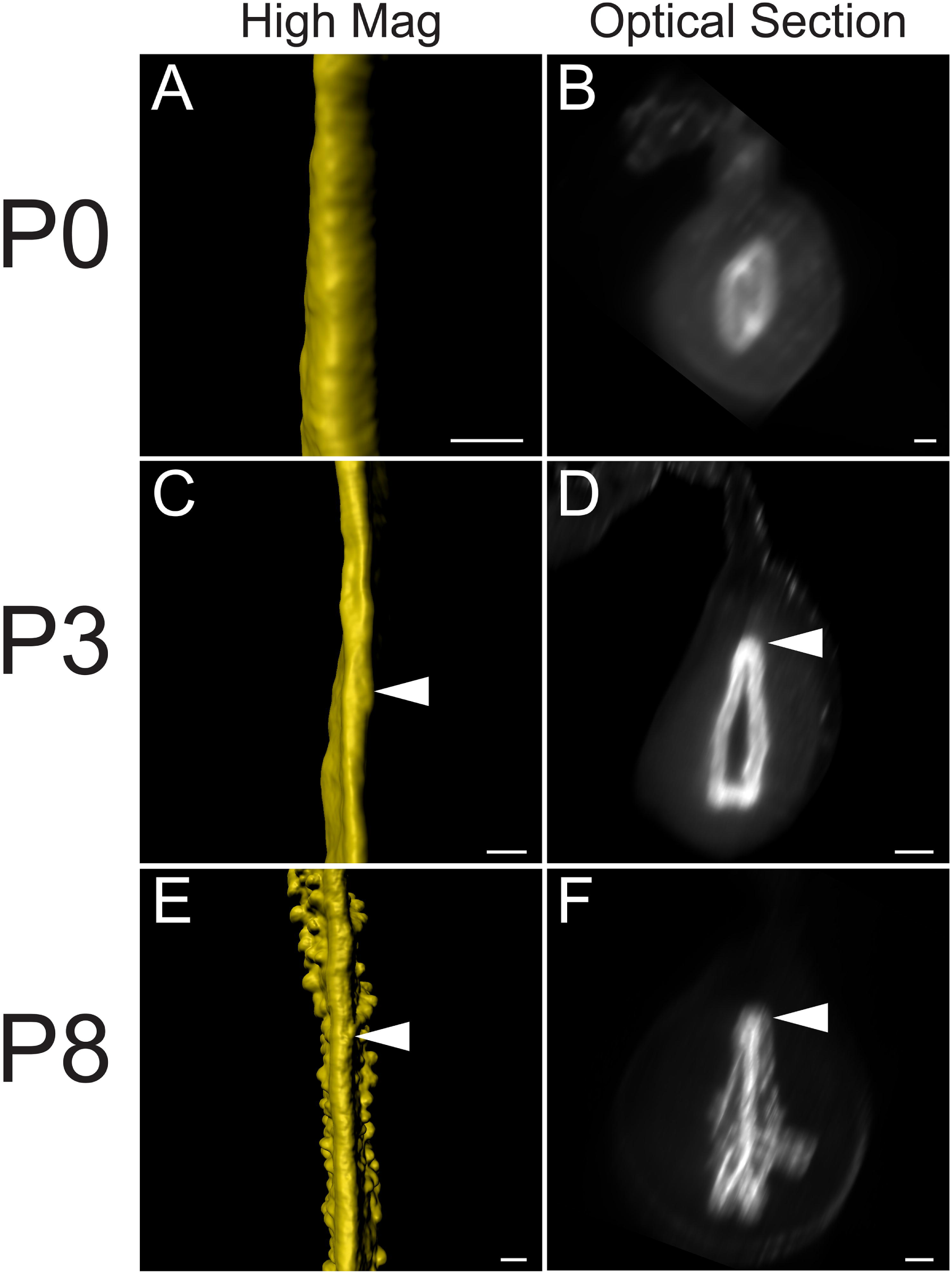
Initial formation of the uterine rail during perinatal development. 3D reconstructions of the perinatal epithelium, highlighting the uterine rail. (**A, C, E**) High magnifications of pseudo-colored surface renderings of the uterine epithelium. Mesometrial view. (**B, D, F**). Optical cross-sections of light sheet images containing fluorescent TROMA-1 stained uterine epithelium. Mesometrium to the top. Arrowheads correspond to the uterine rail. (**A-B**) P0, (**C-D**) P3, and (**E-F**) P8. Scale bars for (**A-F**) = 100 μm.

### Timing of gland formation in the perinatal mouse uterus

Uterine glands are thought to be derived from the luminal epithelium and are contiguous with the luminal epithelium as they mature (Gray, Bartol et al. 2001). Glands are observed as discrete round/oval epithelial units in the stromal compartment in histological sections (Filant, Zhou et al. 2012). Previous studies have suggested that, uterine glands form at P5 (Cooke, Ekman et al. 2012). To determine when the uterine epithelium first form buds (Stage 1, Vue et al., 2018) during adenogenesis, we examined the uterine epithelium before P5. At P3, there was no evidence of adenogenesis (**Fig. 3A-C**, n=6). At P4, the beginning stages of epithelial bud formation were apparent as the epithelium had begun to invade the stroma (**Fig. 3D-F,** n=6). At P5, the epithelial buds are more pronounced and rounded (**Fig. 3G-I**, n=7). At P4 and P5, the bud-stage glands were homogeneously distributed throughout the uterine epithelium.

**Fig. 3.**
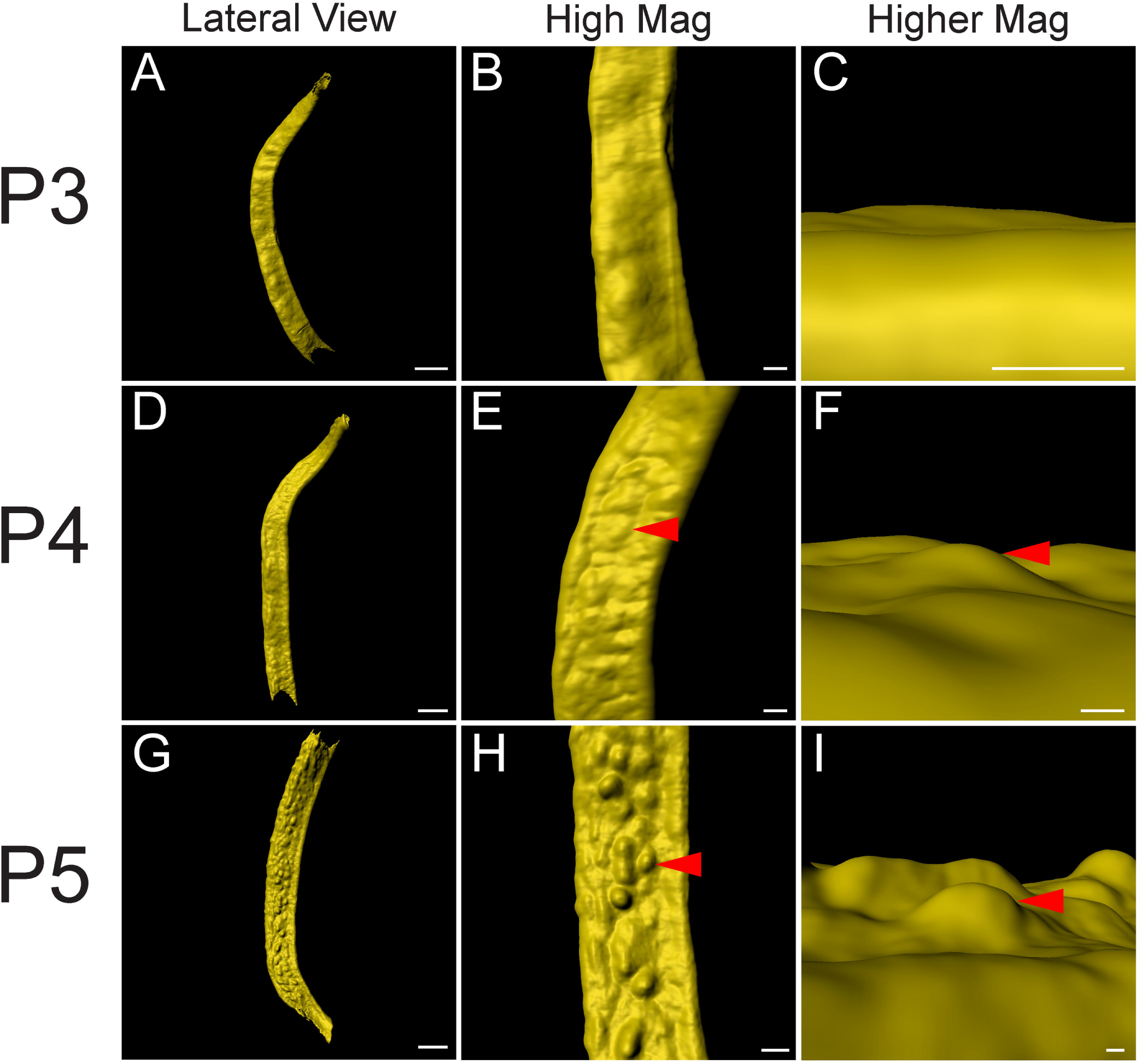
Initial formation of uterine glands in the epithelium. Initial formation of epithelial invaginations in the perinatal mouse uterus. (**A, D, G**) Pseudo-colored surface renderings of the uterine epithelium. Lateral view. (**B, E, H**) High magnification surface renderings of the uterine epithelium corresponding to the first column. Lateral view. (**C, F, I**) Higher magnification of surface rendered single forming glands corresponding to second column. Mesometrial view. Red arrowheads point to forming glands. (**A-C**) P3, (**D-F**) P4, and (**G-I**) P5. Scale bars for (**A, D, G**) = 400 μm. Scale bars for (**B, E, H**) = 100 μm. Scale bars for (**C, F, I**) = 50 μm.

### Identification of a ventral epithelial ridge, a novel structure in the perinatal mouse uterine horn

During our 3D imaging of the perinatal uterus we discovered another novel epithelial structure that runs along the antimesometrial pole of each uterine horn (**Fig. 4**). At P6-P8, we found a ridge that runs along the longitudinal axis. Along the ridge there are two rows of forming glands at the bud stage (Stage 1) (**Fig. 4E-J**). Thus, we have termed this the “ventral ridge.” The ventral ridge is another indication that dorsoventral patterning is occurring in the perinatal uterus. This distinct morphological structure, along with the two rows of glands, can be observed by P6 (**Fig. 4E, F**). However, the first sign of ventral ridge formation can be seen as early as P4 at the mid-anterior portion of the uterine horn (**Fig. 4A, B**). At P5, the ventral ridge is also found at the anterior portion of the uterine horn (**Fig. 4C, D**). At P6, the ventral ridge is fully formed along the entire length of the anteroposterior axis (**Fig. 4E, F**) and becomes more pronounced as the mouse ages at P7 and P8 (**Fig. 4G-J**).

**Fig. 4.**
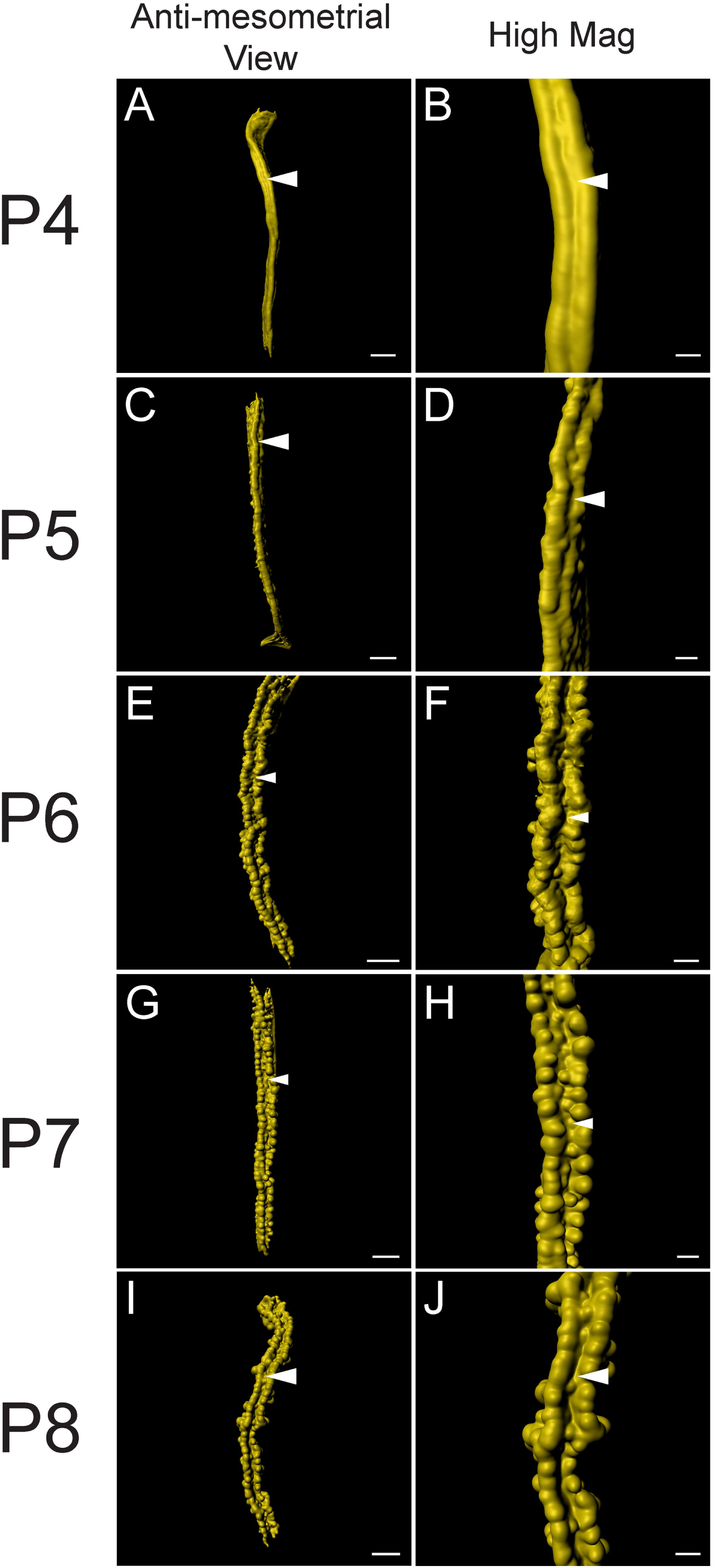
Identification of the ventral ridge in the perinatal mouse uterus. 3D reconstructions of TROMA-1 stained perinatal mouse uterine epithelium. (**A, C, E, G, I**) Pseudo-colored surface renderings of the uterine epithelium. Anti-mesometrial view. (**B, D, F, H, J**) High magnification surface renderings of the uterine epithelium corresponding to the first column. Anti-mesometrial view. White arrowheads, ventral ridge. (**A-B**) P4, (**C-D**) P5, (**E-F**) P6, (**G-H**) P7, and (**I-J**) P8. Scale bars for (**A, C, E, G, I**) = 400 μm. Scale bars for (**B, D, F, H, J**) = 150 μm.

### Lateral epithelial folds mark segments along the anteroposterior axis of the perinatal uterine horn

By P8, prominent anterior to posterior epithelial folds were observed, extending laterally from the lumen into the stroma (**Fig. 5**). When the epithelium is visualized as a surface rendering, these folds appear to be spaced relatively evenly along the anteroposterior axis of the uterine horn. We have termed these structures “uterine folds” and the region between the folds we term “uterine segments” (**Fig. 5M-P**). We find 3-4 glands extend from each uterine fold. The uterine folds can be recognized at P6 (**Fig. 5F-H**) and P7 (**Fig. J-L**), and P8 (N-P). These folds and segments are not apparent at P5 (**Fig. 5A-D**). These observations suggest that uterine folds form between P5 and P6.

**Fig. 5.**
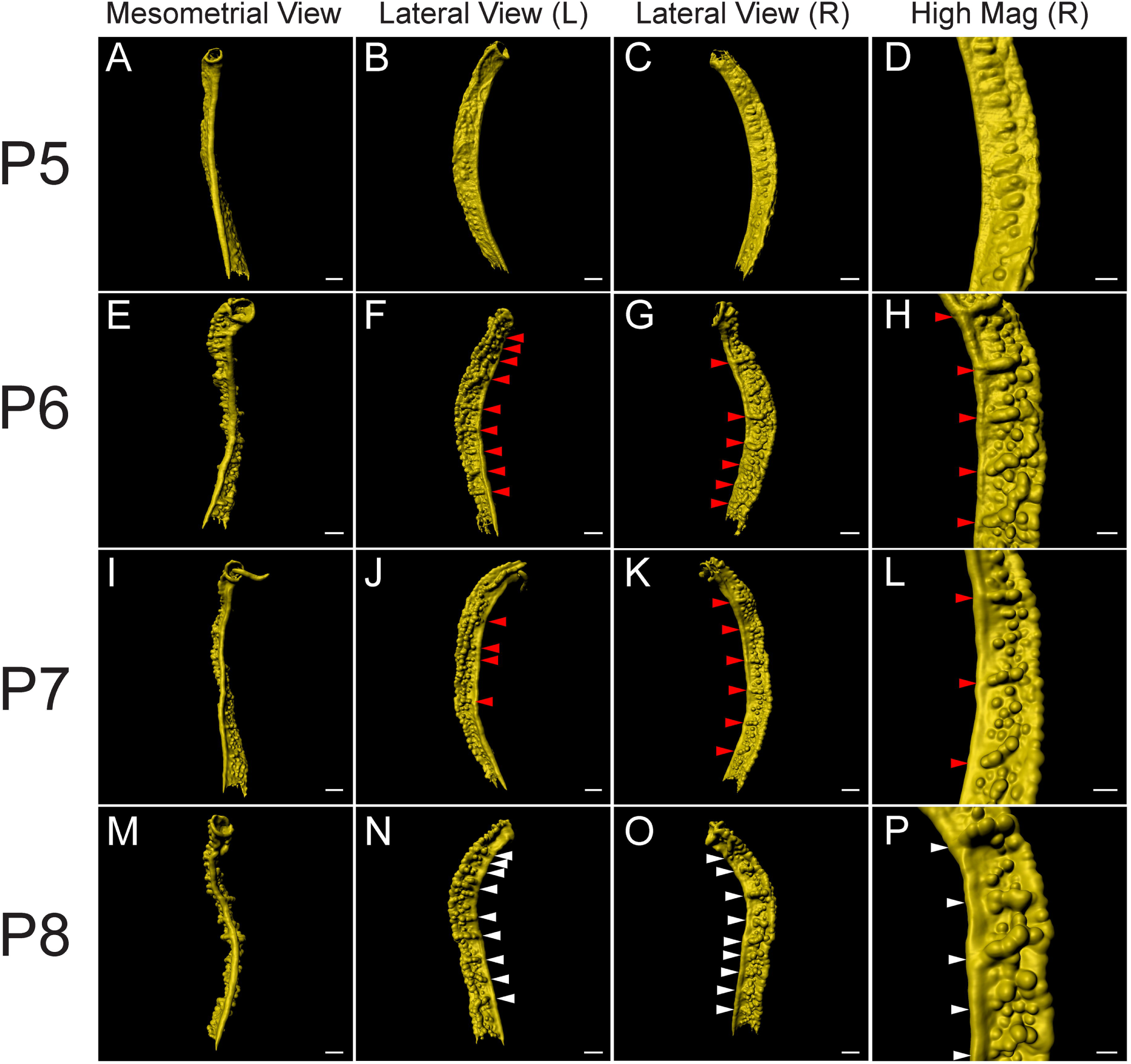
Identification of lateral segments along the anteroposterior axis of the perinatal mouse uterus. 3D reconstructions of the mouse uterine epithelium, highlighting lateral segments along the anteroposterior axis. (**A, E, I, M**) Pseudo-colored surface renderings of the uterine epithelium. Mesometrial view. (**B, F, J, N**) Pseudo-colored surface renderings of the uterine epithelium corresponding to the first column. Left lateral view. (**C, G, K, O**) Pseudo-colored surface renderings of the uterine epithelium corresponding to the first column. Right lateral view. (**D, H, L, P**) High magnification surface renderings of the uterine epithelium corresponding to the third column. Red arrowheads correspond to forming lateral epithelial folds. White arrowheads correspond to the lateral epithelial folds. (**A-D**) P5, (**E-H**) P6, (**I-L**) P7, and (**M-P**) P8. Scale bars for (**A-C, E-G, I-K, M-O)** = 300 μm. Scale bars for (**D, H, L, P**) = 150 μm.

We examined the locations of the epithelial folds at P8.The uterine folds appear to be present in an alternating left-right pattern, relative to the uterine rail (**Fig. 6A-D**). This creates a zigzag-like morphology (**Fig. 6C**). We measured the distance between the uterine folds, i.e. the length of uterine segments. We found that the average distance between uterine folds (n=14) is 317.9 ± 61.9 um (**Fig. 6E**). However, most of the uterine segments were approximately 200 um in length.

**Fig. 6.**
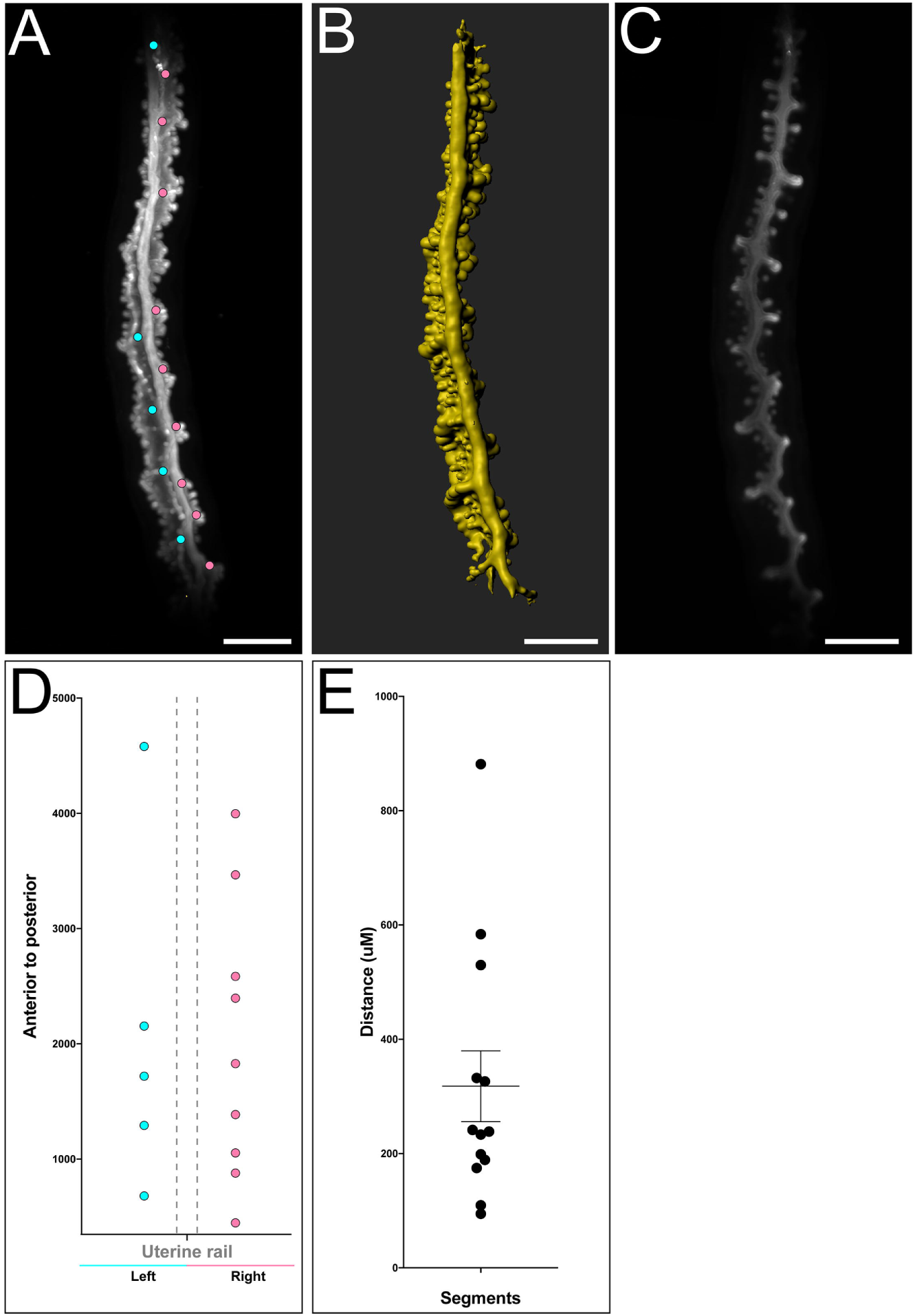
Distribution of lateral epithelial folds in the perinatal mouse uterus. Uterine lateral segment length was measured at P8 using Imaris FilamentsTracer. (**A**). Raw maximum projection of the mouse epithelium. Each dot represents the location of a lateral epithelial fold along the anteroposterior axis. A blue dot represents a lateral epithelial fold on the left side and a pink dot represents a lateral epithelial fold on the right side, relative to the uterine rail, mesometrial view. (**B**) Pseudo-colored surface rendering of the uterine epithelium corresponding to the first column, mesometrial view. (**C**). Single crosssection of the uterine epithelium corresponding to the first column. (**D**). Location (microns) of lateral epithelial folds along the anteroposterior axis for the specimen shown in **A**. (**E**). Average distance between each lateral segment. Error bars indicate average distance ± standard error mean.

## Discussion

The endometrium of the adult uterus is an essential tissue for embryo implantation, growth, and development. The endometrium is composed of luminal and glandular epithelial and stromal tissue compartments surrounded by two layers of smooth muscle comprising the myometrium. The current study focuses on the morphogenetic changes that occur in the epithelial compartment of the perinatal mouse uterus. We used light sheet fluorescent microscopy to generate 3D volumetric reconstructions of the epithelium of the perinatal mouse uterus (Vue et al., 2018). We describe multiple morphological changes in the perinatal uterine epithelium, leading to asymmetries, including the timing of the formation of the uterine rail, the initiation of adenogenesis, a novel structure we term the ventral ridge, and lateral epithelial folds alternating left and right that appear to define uterine segments along the anteroposterior axis.

The mouse uterine horns are suspended in the peritoneal cavity by the broad ligament. The broad ligament attaches to the mesometrial side of the uterine horn, defining a mesometrial-antimesometrial axis. This mesometrial-antimesometrial axis can be considered a dorsal-ventral axis. The lateral and ventral regions of the adult mouse uterus have endometrial or uterine glands, whereas the dorsal region lacks these glands (Arora et al., 2016; Yuan et al., 2017; Vue et al., 2018).

Patterning of the female reproductive tract has been studied extensively along the anteroposterior axis of the Müllerian duct, which results in its segmentation into structurally and functionally unique regions of the female reproductive tract that includes the oviducts, uterus, cervix and anterior region of the vagina (Kobayashi and Behringer, 2003). Within the uterus, radial patterning begins during embryogenesis and subsequently after birth establishes the endometrium, myometrium and perimetrium (Bartol et al. 1993, Bartol et al. 1999, Gray et al. 2001).

At birth, the lumen of the uterus is lined by the luminal epithelium with no morphological dorsal-ventral polarity although anatomically the presence of the mesometrium defines a dorsal-ventral axis for the organ. Thus, dorsal-ventral epithelial patterning occurs after birth (Goad et al., 2017; Vue et al., 2018). Recently, we used light sheet fluorescent microscopy and 3D reconstructions to identify a dorsal epithelial structure in the postanatal (P8, P11, and P21) uterus we termed the uterine rail (Vue et al., 2018). Our current 3D reconstructions of the perinatal mouse uterus showed the initial formation of the uterine rail at P3. The uterine rail becomes more prominent each subsequent day between P4 and P8. Thus, the perinatal uterine epithelium acquires dorsal-ventral pattern by P3. Dorsally-restricted Wnt activity has been observed in the postnatal mouse uterus using *TCF-GFP* WNT-reporter transgenic mice (Goad et al., 2017). At P3, GFP expression was observed throughout the uterine epithelium. However, at P6, GFP expression was up-regulated in the dorsal uterine epithelium, suggesting a localization of WNT-regulated gene expression in the dorsal uterine epithelium at P6. However, the earlier timing of the initial formation of the uterine rail suggests that its development may be WNT-independent.

Uterine glands and their secretions are essential for implantation (Filant and Spencer, 2013). Previous studies and our current results show that no glands are present at P3 (Hu et al. 2004, Goad et al. 2017). Forming uterine glands have been identified at P5 in multiple different genetic backgrounds (Cooke et al. 2012). We found Stage 1 (bud stage) developing uterine glands at P5 consistent with previous observations. Our 3D reconstructions at P4 show that initial buds are already present. Thus, we conclude that initial gland formation occurs between P3 and P4. Defining the precise timing for the initiation of uterine gland formation should facilitate the search for molecular regulators of adenogenesis from developmental transcriptome datasets (Filant and Spencer, 2013; Wu et al., 2017).

During our volumetric analysis of the perinatal mouse uterine epithelium we discovered a ventral structure composed of two rows of glands. These two parallel rows of Stage 1 glands (bud stage, Vue et al., 2018) were readily found at P6 to P8 but indications of a ventral structure were apparent even at P4. We have named this structure the “ventral ridge”. The 3D rendering of the perinatal epithelium revealed this structure that was not apparent in previous 2D histology (Hu et al., 2004). DICKKOPF proteins are secreted negative regulators of WNT activity (Niehrs, 2006). *Dkk2* transcripts were found to be localized in the anti-mesometrial stroma of the P6 and P15 mouse uterus (Goad et al., 2017). Perhaps an inhibition of WNT activity in the antimesometrial (ventral) stroma creates a permissive environment for the robust development of the ventral ridge that initiated formation at P4. The presence of two parallel rows of forming glands flanking the ventral midline of the uterine epithelium suggests the establishment of a bilateral symmetry.

The adult female mouse uterine horn has anteroposterior and dorsal-ventral pattern. There are anterior to posterior segments, including the oviduct, uterus, cervix, and vagina. Within the uterus, there is dorsal to ventral pattern, including the dorsally-positioned uterine rail, lateral/ventral endometrial glands, and now in the perinatal uterus, the ventral ridge (Hu et al., 2004; Vue et al., 2018). Our volumetric analysis also showed expansion of luminal epithelial folds that mark segmental patterning along the anteroposterior axis, suggesting a form of segmentation within the developing perinatal uterine horn. The presence of luminal epithelial folds was found at P8 although there were morphological indications of segments as early as P6. Interestingly, the luminal epithelial folds were not bilaterally symmetrical but rather appeared to alternate in a left-right pattern, that at later stages correlated with a zigzag-like morphology when viewed dorsally. Conditional loss of the tumor suppressor gene *Nf2* in the uterine epithelium results in a lack of uterine glands (Lopez et al., 2018). Interestingly, at P7 there were lateral epithelial folds that alternated in a left-right pattern. These results suggest that the lateral epithelial folds are independent of gland formation

In summary, light sheet fluorescent microscopy is a powerful imaging tool to render 3D volumetric reconstructions, revealing diverse morphological changes that occur in the epithelium of the perinatal mouse uterus. Our studies define a set of morphogenetic patterns that develop soon after birth in the mouse uterine epithelium that ultimately result in the formation of the mature endometrium that is required for reproduction

## Experimental Procedures

### Mice

Swiss Webster outbred mice (Taconic Biosciences) were used in this study. All mice were maintained in compliance with the Public Health Service Policy on Humane Care and Use of Laboratory Animals, the U. S. Department of Health and Human Services Guide for the Care and Use of Laboratory Animals, and the United States Department of Agriculture Animal Welfare Act. All protocols were approved by the University of Texas MD Anderson Cancer Center Institutional Animal Care and Use Committee.

### Immunostaining, image capture and post-acquisition processing

Dissections and immunostaining were performed as described (Vue, Gonzalez et al. 2018). Antibody used in this study for all images was TROMA-1 (Developmental Studies Hybridoma Bank, Iowa USA, 1:100). Whole-mount fluorescent images were obtained using a Zeiss Lightsheet Z.1 Light Sheet Fluorescence Microscope (LSFM) (Zeiss, Jena Germany) with 405/488/561/640 nm lasers. A 5×, NA0.6 Plan-Apochromat water immersion objective (Carl Zeiss) and dual-sided illumination were used for imaging. Laser power and exposure times varied depending on the amount of fluorescent signal and proximity of the signal to the surface of the tissue. All Z-stacks were processed by default settings through Imaris (Bitplane), including pseudo-coloring, adjusting the dynamic range of each color channel, surface rendering, smoothing with surface detail and background subtraction. FilamentTracer software (Imaris, Bitplane), a program usually used to detect neurons, microtubules and other filament-like structures, was adapted to determine the length of luminal segments. Each luminal segment (nearest the uterine rail) was manually demarcated as the base of a “dendrite” by using default settings. From this, the location of lateral epithelial folds were determined and intersegment distance quantified.

### Statistical Analysis

Statistical analysis was performed using the Prism 6.0 software (GraphPad). The length between uterine folds was determined by performing column analyses and column statistics for all samples. For statistical analyses, α=0.05 was used to determine significance.

## Acknowledgements

We thank Rachel D. Mullen and Shuo-Ting Yen for helpful discussions. We thank Chih-Wei Hsu, Tegy John Vadakkan and Jason M. Kirk from the Baylor College of Medicine Optical Imaging and Vital Microscopy Core for microscopy and assistance, Adriana Paulucci and Henry Adams from the MD Anderson Department of Genetics Microscopy Core for microscopy, assistance and post-image acquisition processing, Kenneth Trimmer for his help with the Imaris software, and the IDDRC Microscopy Core for their help with Imaris Filaments.

## Movie legends

**Movie 1**. TROMA-1 whole-mount immunofluorescence and surface-rendered opaque view of P3 uterine horn.

**Movie 2**. TROMA-1 whole-mount immunofluorescence and surface-rendered opaque view of P4 uterine horn.

**Movie 3**. TROMA-1 whole-mount immunofluorescence and surface-rendered opaque view of P5 uterine horn.

**Movie 4**. TROMA-1 whole-mount immunofluorescence and surface-rendered opaque view of P6 uterine horn.

**Movie 5**. TROMA-1 whole-mount immunofluorescence and surface-rendered opaque view of P7 uterine horn.

**Movie 6**. TROMA-1 whole-mount immunofluorescence and surface-rendered opaque view of P8 uterine horn.

## References

Arora, R., A. Fries, K. Oelerich, K. Marchuk, K. Sabeur, L. C. Giudice and D. J. Laird (2016). “Insights from imaging the implanting embryo and the uterine environment in three dimensions.” Development 143(24): 4749–4754.

Bartol, F. F., A. A. Wiley, J. G. Floyd, T. L. Ott, F. W. Bazer, C. A. Gray and T. E. Spencer (1999). “Uterine differentiation as a foundation for subsequent fertility.” J Reprod Fertil Suppl 54: 287–302.

Bartol, F. F., A. A. Wiley, T. E. Spencer, J. L. Vallet and R. K. Christenson (1993). “Early uterine development in pigs.” J Reprod Fertil Suppl 48: 99–116.

Belle, M., D. Godefroy, G. Couly, S. A. Malone, F. Collier, P. Giacobini and A. Chedotal (2017). “Tridimensional Visualization and Analysis of Early Human Development.” Cell 169(1): 161–173.e112.

Branham, W. S., D. M. Sheehan, D. R. Zehr, E. Ridlon and C. J. Nelson (1985). “The postnatal ontogeny of rat uterine glands and age-related effects of 17 beta-estradiol.” Endocrinology 117(5): 2229–2237.

Brody, J. R. and G. R. Cunha (1989). “Histologic, morphometric, and immunocytochemical analysis of myometrial development in rats and mice: I. Normal development.” Am J Anat 186(1): 1–20.

Brulet, P., C. Babinet, R. Kemler and F. Jacob (1980). “Monoclonal antibodies against trophectoderm-specific markers during mouse blastocyst formation.” Proc Natl Acad Sci U S A 77(7): 4113–4117.

Cooke, P. S., G. C. Ekman, J. Kaur, J. Davila, I. C. Bagchi, S. G. Clark, P. J. Dziuk, K. Hayashi and F. F. Bartol (2012). “Brief exposure to progesterone during a critical neonatal window prevents uterine gland formation in mice.” Biol Reprod 86(3): 63.

Filant, J. and T. E. Spencer (2013). “Endometrial glands are essential for blastocyst implantation and decidualization in the mouse uterus.” Biol Reprod 88(4): 93.

Filant, J., H. Zhou and T. E. Spencer (2012). “Progesterone inhibits uterine gland development in the neonatal mouse uterus.” Biol Reprod 86(5): 146, 141–149.

Goad, J., Y. A. Ko, M. Kumar, S. M. Syed and P. S. Tanwar (2017). “Differential Wnt signaling activity limits epithelial gland development to the anti-mesometrial side of the mouse uterus.” Dev Biol 423(2): 138–151.

Gray, C. A., F. F. Bartol, B. J. Tarleton, A. A. Wiley, G. A. Johnson, F. W. Bazer and T. E. Spencer (2001). “Developmental biology of uterine glands.” Biol Reprod 65(5): 1311–1323.

Hondo, E., T. Phichitrasilp, K. Kokubu, K. Kusakabe, N. Nakamuta, H. Oniki and Y. Kiso (2007). “Distribution patterns of uterine glands and embryo spacing in the mouse.” Anat Histol Embryol 36(2): 157–159.

Hu, J., C. A. Gray and T. E. Spencer (2004). “Gene expression profiling of neonatal mouse uterine development.” Biol Reprod 70(6): 1870–1876.

Kurita, T., P. S. Cooke and G. R. Cunha (2001). “Epithelial-stromal tissue interaction in paramesonephric (Mullerian) epithelial differentiation.” Dev Biol 240(1): 194–211.

Kobayashi, A. and R. R. Behringer (2003). “ Developmental genetics of the female reproductive tract in mammals.” Nat Rev Genet 4(12): 969–980.

Lopez, E. W., Z. Vue, R. R. Broaddus, R. R. Behringer and A. B. Gladden (2018). “The ERM family member Merlin is required for endometrial gland morphogenesis.” Dev Biol 442(2): 301–314.

Niehrs C. (2006). Function and biological roles of the Dickkopf family of Wnt modulators. Oncogene 25(57): 7469–7481.

Vue, Z., G. Gonzalez, C. A. Stewart, S. Mehra and R. R. Behringer (2018). “Volumetric imaging of the developing prepubertal mouse uterine epithelium using light sheet microscopy.” Mol Reprod Dev 85(5): 397–405.

Wu B., C. An, Y. Li, Z. Yin, L. Gong, Z. Li, Y. Liu, B. C. Heng, D. Zhang, H. Ouyang and X. Zou. (2017). **”**Reconstructing Lineage Hierarchies of Mouse Uterus Epithelial Development Using Single-Cell Analysis.**”** Stem Cell Rep 9(1): 381–396. (2017).

Yuan, J., W. Deng, J. Cha, X. Sun, J. P. Borg and S. K. Dey (2018). “Tridimensional visualization reveals direct communication between the embryo and glands critical for implantation.” Nat Commun 9(1): 603.

